# Human short association fibers are thinner and less myelinated than long fibers

**DOI:** 10.1101/2024.10.21.619354

**Authors:** Philip Ruthig, David Edler v.d. Planitz, Maria Morozova, Katja Reimann, Carsten Jäger, Tilo Reinert, Siawoosh Mohammadi, Nikolaus Weiskopf, Evgeniya Kirilina, Markus Morawski

## Abstract

The size and complexity of the human brain requires optimally sized and myelinated fibers. White matter fibers facilitate fast communication between distant areas, but also connect adjacent cortical regions via short association fibers. The fundamental questions of i) how thick these fibers are and ii) how strongly they are myelinated, however, remain unanswered. We present a comprehensive analysis of ∼400,000 fibers of human white matter regions with long (corpus callosum) and short fibers (superficial white matter). We demonstrate a substantially smaller fiber diameter and lower myelination in superficial white matter than in the corpus callosum. Surprisingly, we do not find a difference in the ratio between axon diameter and myelin thickness (g-ratio), which is close to the theoretically optimal value of ∼0.6 in both areas. For the first time, to our knowledge, we shed light on a fundamental principle of brain organization that will be essential to understand the human brain.

## Introduction

White matter connects neighboring and distant cortical areas. As such, white matter is crucial for various functions, e.g. in sensory-motor integration (1,2), language perception (3), hemispheric information transfer (4), and its plasticity is highly behaviorally relevant, e.g. during learning (5). Reflecting these various functional roles, white matter fibers can be highly variable in length and diameters (6). Long connections include corticospinal projections of up to 1m length (7,8) and projections to the prefrontal cortex (9), while short connections of only a few centimeters length can be found between areas such as the motor and somatosensory cortex (1,10,11), different visual cortical areas (12), or auditory cortex fields (13).

Axon radius and myelination thickness are the main factors defining conduction speed of a fiber, but at the same time also defining the fibers energy demand and the space it occupies. The higher in diameter an axon and its myelination is, the faster it can propagate signals between distant regions of the human brain. On the flipside, increasing fiber and myelination thickness also increases the metabolic upkeep of a fiber (even when not firing) dramatically (14,15), and increases the required space. Therefore, only very few axons have a high diameter and thick myelination (16).

Theoretically, a fiber optimized for maximum conduction speed in the CNS has a g-ratio (ratio of axon diameter to outer fiber diameter) of 0.6 to 0.77 (17,18). In an evolutionary sense, the tradeoff for thicker and more myelinated fibers is only positive when a functional reason for fast and efficient transmission of signals exists. Such functional reasons might include rapid integration of somatosensory feedback to the motor cortex and vice versa (11), relaying temporally complex auditory cues to higher order areas (19–21), or rapid conduction of visual stimuli via the optic nerve to the visual cortex (22). A long-standing hypothesis in the field is that the longer a fiber tract is, the larger is the axon diameter and the thicker is the myelination of the local fiber population, as this allows more efficient and faster information transfer over long distances (23–26). Although this hypothesis is widely spread, to date there exists no ultrastructural, gold standard data from human brains to support this hypothesis.

In this study, we provide insight into fundamental principles of white matter organization. We investigate structural properties of axons located in regions with either predominantly short or predominantly long range fibers. We compared two sets of regions using transmission electron microscopy: (i) long range fibers sampled from the entire corpus callosum (CC), (ii) short range fibers sampled in superficial white matter (SWM, sampled <3mm from the white/gray matter border) between primary motor (M1) and somatosensory cortex (S1), and between the primary (V1) and secondary (V2) visual cortex (Fig. 1). In the chosen SWM regions, we expect a higher percentage of short association fibers connecting M1 and S1 and V1 and V2 by so-called ‘U-fibers’. These short fiber tracts have been documented perpendicular to the central sulcus (1,10,11), and in the visual cortex (12,27), albeit only with gross anatomical methods and diffusion MRI. A detailed ultrastructural description is lacking. The closest analogue study, to our knowledge, was conducted in fetal sheep brain and shows that SWM has 50% more thin axons (diameter <0.65 µm) than the CC, and slightly more thick (diameter >8.5 µm) axons (28).

Previous studies on the topic have either employed relatively distant model organisms or relied on exceedingly small fields of views and number of manually segmented fibers (28–30). In the present study, we employ a state of the art automated segmentation algorithm to analyze high quality human material, showing that short association fibers have smaller diameters compared to long-range fibers but have a similar mean g-ratio, close to the optimal value of ∼0.6. For the first time, to our knowledge, we provide experimental evidence for the distinct structural properties of short association fibers in humans, highlighting the tight link between function and structure in human white matter.

**Fig. 1:**
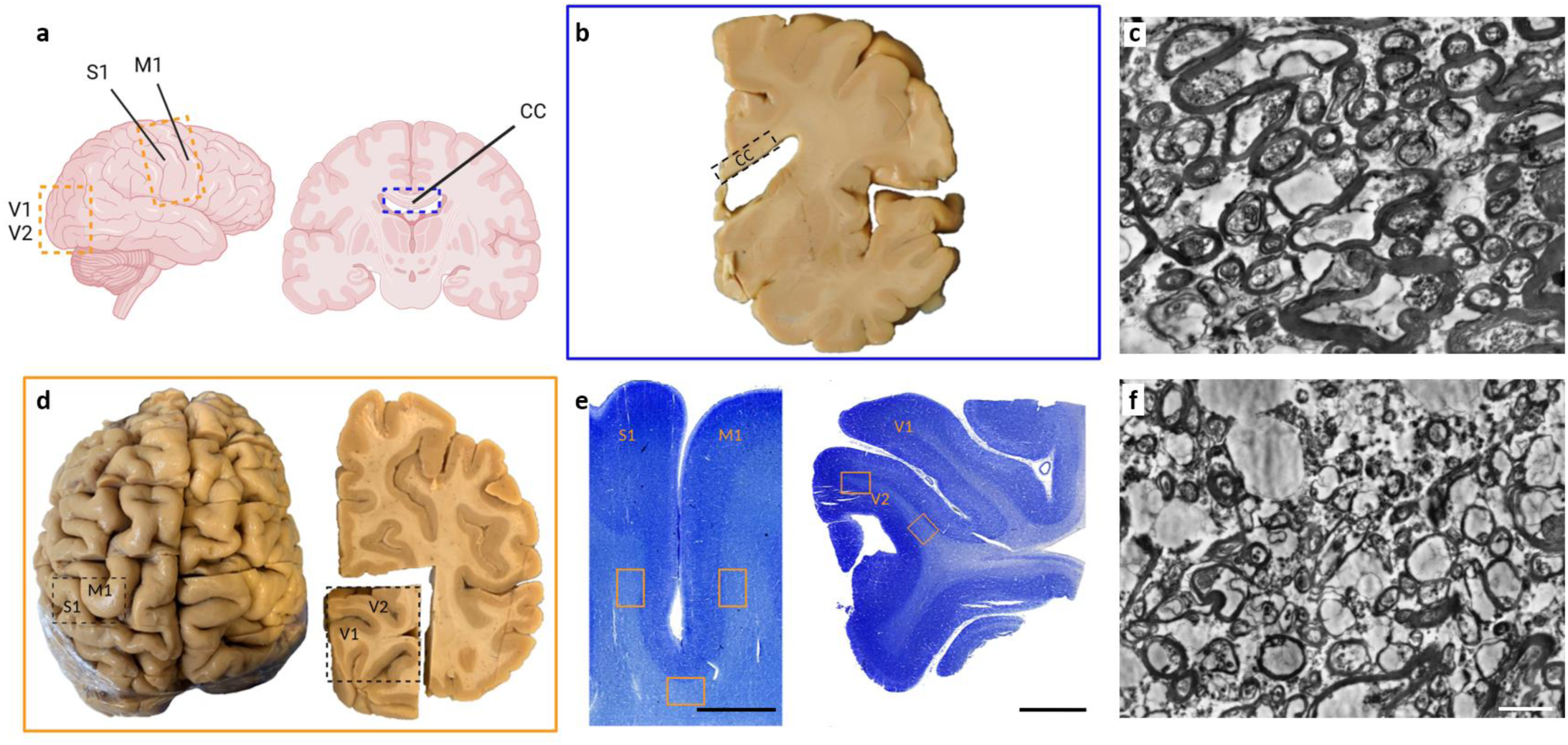
Overview over brain regions sampled in this study. **a**, Depiction of a human brain and brain slice with used SWM regions (yellow) and CC (blue) highlighted. **b**, Gross anatomical view of coronal slice, containing CC. **c**, Transmission electron micrograph of CC tissue. **d**, gross anatomical location of SWM tissue. **e**, Nissl stainings of SWM tissue along the central sulcus (left) and visual cortex (right), with example regions that were used highlighted (yellow). Scale bars are 5 mm. **f**, Transmission electron micrograph of SWM tissue. Scale bar is 2 µm.

## Results

### Short fibers have smaller axon diameters and thinner myelin as compared to long fibers

To understand systematic differences between long and short fibers, we assessed the outer fiber diameter (axon + myelination diameter) and axon diameter (here assumed to be identical with the inner myelin diameter) in SWM and CC. In contrast to previous studies, which frequently only sampled few fibers, we automatically analyzed ∼200 000 fibers in CC and SWM each. We generated samples from 5 donor brains (4 male, 1 female, aged 60-78 years) and generated the data with transmission electron microscopy (Fig. 2a). To reliably segment myelinated fibers, we trained a variant of a U-Net (DenseNet, (31)), which outputs a per-pixel prediction of the raw data (Fig. 2b). This preliminary segmentation is then processed to a final segmentation of each axon + myelin pair (Fig. 2c), which is then measured and analyzed (Fig. 2d-f). We show that axons in SWM are systematically thinner (mean axon diameter of 0.57 µm (SWM) vs 0.75 µm (CC)) and less myelinated than in CC (mean myelin diameter: 1.24 µm (SWM) vs 1.57 µm (CC), see Fig. 3a, b). We describe these distributions by fitting a Generalized Extreme Value function (GEV) to each distribution of axon diameters and outer fiber diameters using Markov chain Monte Carlo sampling (Fig. 3a, Fig. 4, Fig. S7, S8). Each GEV is defined by a location parameter µ (defining the location of the distribution along the x axis), scale parameter σ (defining the size of the distribution), and shape parameter ξ (defining the tailedness of the distribution). We display these parameters in Fig. 4b, showing that the distribution for CC axons and myelination is more tailed towards large diameters and has a higher location parameter than the GEV corresponding to SWM. Furthermore, the scale parameter of both axon and outer fiber diameter distribution is higher in CC.

**Fig. 2:**
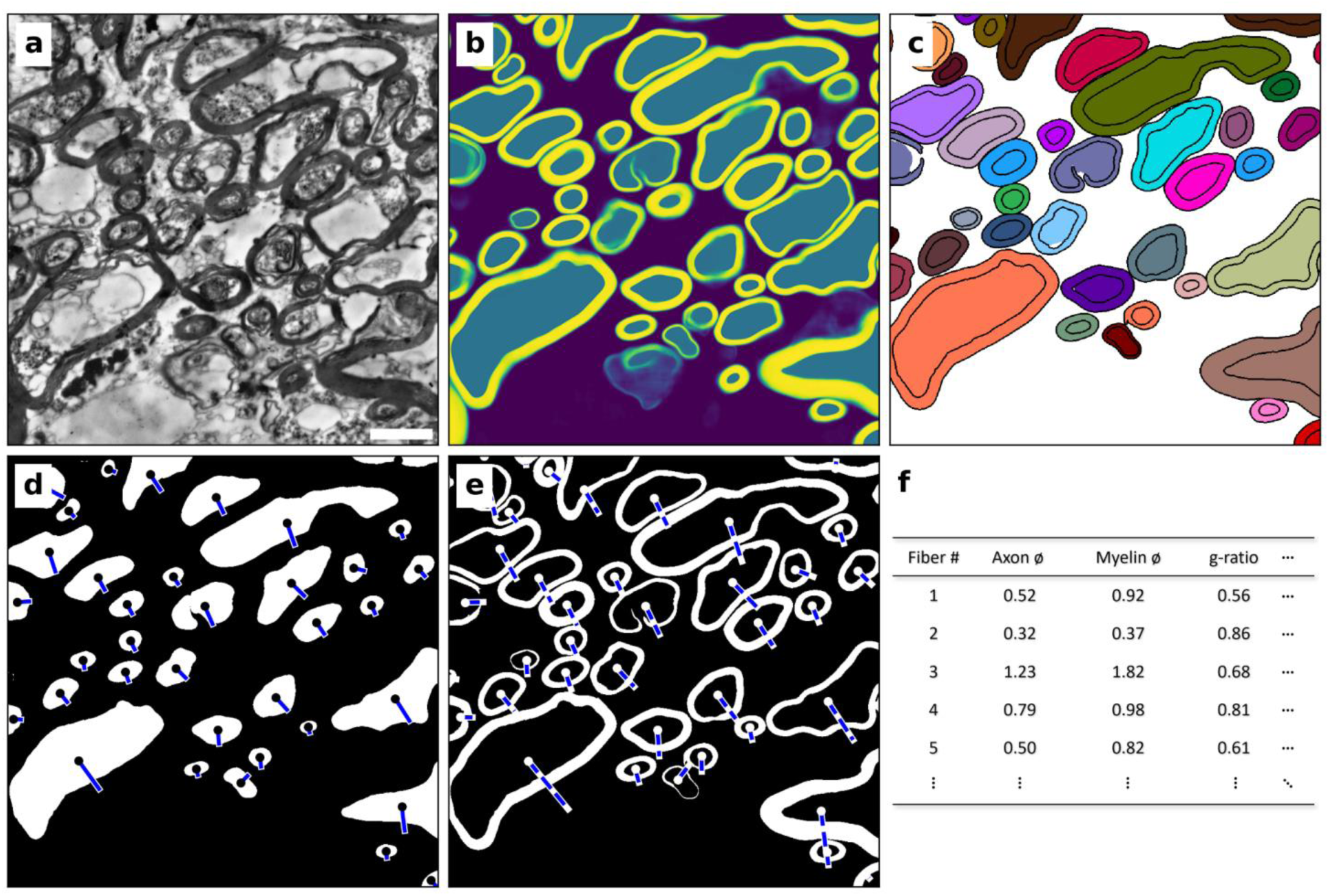
Overview of the segmentation approach for electron microscopic data. **a**, Preprocessed transmission electron microscopic data of human CC. **b**, Semantic prediction of the preprocessed data by the trained DenseNet, returning a likelihood of each pixel to belong to either background (violet), axon (teal), or myelin (yellow). **c**, Post-processed final segmentation. Each instance of a fiber (axon and myelin sheath) is labelled with a different random color. **d,e**, Measurement of axon (**d**) and myelination (**e**) diameters based on ellipsoids fitted to each structure. Points show the centroid of each structure, axes show half of the minor axis of the fitted ellipse for the axon (solid line) and myelin (dashed line). **f**, Measurements (SWM: n = 187307, CC: n = 203688) from d, e are used for statistical assessment of local ultrastructure. We treat the short axis (red) of the fitted ellipse as the actual diameter of each structure. Scale bar = 2 µm.

### Superficial white matter fibers are more diverse than Corpus Callosum fibers

But is this difference also reflected in the g-ratio? We find that, although CC and SWM are very different in axon and outer fiber diameter (Fig. 3a,b), the mean g-ratio of both groups is almost identical (SWM: 0.511, CC: 0.505, Fig. 3e). In contrast, the shape of the g-ratio distribution shows a higher variety of different axon/myelin thickness combinations in SWM than in CC. This is also reflected in the relationship of axon diameter to g-ratio (Fig. 3c, d), where larger CC fibers have a higher tendency to cluster around g-ratio 0.6 than SWM fibers.

**Fig. 3:**
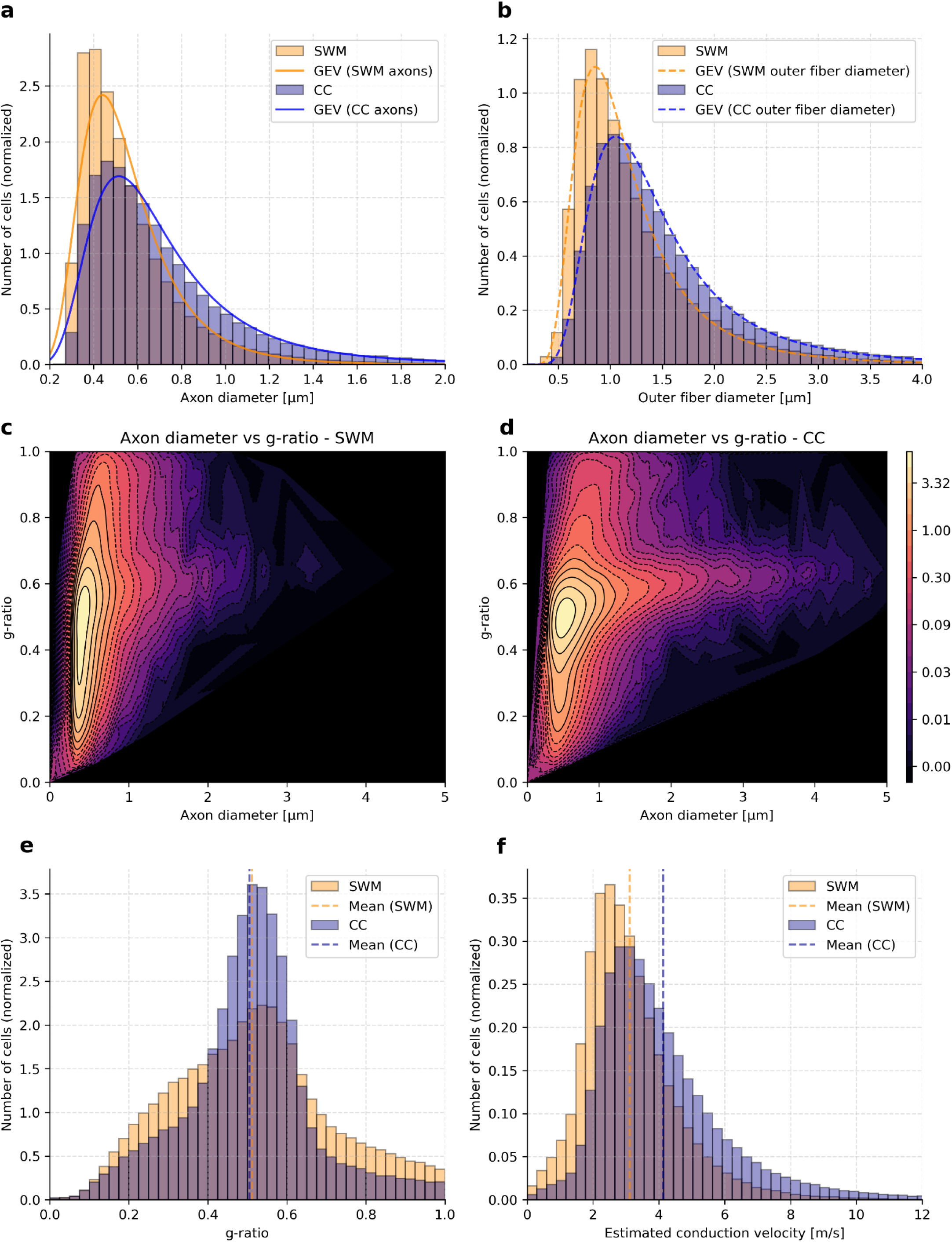
Longer white matter tracts show higher axon diameters, thicker myelination, higher estimated conduction velocities, but no difference in mean g-ratio. **a**,**b** Distributions of SWM & CC axon diameters (a) and outer fiber diameters (b) with fitted Generalized extreme value functions (GEV). **c,d**, Densities of measured axon diameter vs g-ratio in SWM (**c**) and CC (**d**). Contour levels are estimated from Gaussian kernel density in log scale. **e**, Histogram of g-ratio distribution in CC and SWM. SWM mean is 0.511, CC mean is 0.505. **f**, Estimated conduction velocities of CC and SWM fibers according to the conduction velocity formula by Rushton et al (18), modified according to Schmidt & Knösche (32). Estimated mean connection speeds are 4.1m/s (CC) and 3.1m/s (SWM).

### Specialization is reflected in region-specific conduction velocity changes

Finally, the important question arises as to what functional effects the measured differences may have. The conduction velocity of a fiber can be estimated from the axon diameter and myelination thickness. Based on our measurements, we predict conduction velocities of both groups according to Rushton (18), which we adapted according to Schmidt & Knösche (32). The differences in axon and myelination thickness result in an approximately 32% higher mean conduction velocity (CC: 4.1m/s, SWM: 3.1m/s) in CC fibers, with an especially higher proportion of fast conducting (>6m/s) fibers (Fig. 3f).

**Fig. 4:**
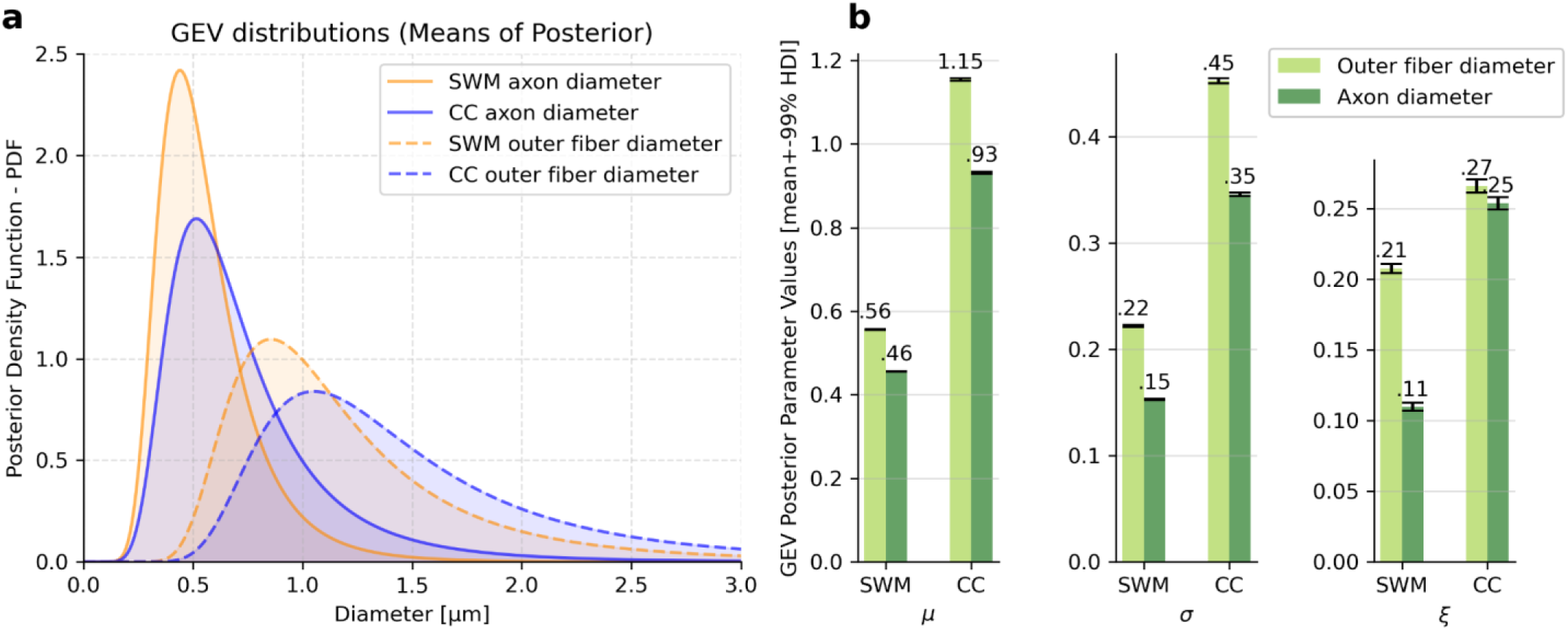
Modeling axon diameter and myelination thickness with Generalized Extreme Value functions. **a**, Generalized extreme value functions (GEV) fitted on SWM & CC axons and outer fiber diameter distributions. Histograms for the underlying data are given in Fig. 2a,b. **b**, Parameters of GEV functions fit to CC and SWM data. Error bars display 99% highest posterior interval range. Displayed parameters are µ (location parameter), σ (scale parameter), and ξ (shape or ‘tailedness’ parameter).

## Discussion

We show robust evidence in favor of the hypothesis that, in humans, short fibers are smaller and less myelinated than long fibers. Our results are supported by a statistical analysis of a large ensemble of ∼400 000 myelinated axons sampled from two SWM and six CC subregions in three human donor brains each and display an approximately 30% higher mean axon and outer fiber diameter in CC compared to SWM (Fig. 3a, b). The only other available study systematically comparing SWM and CC fibers with electron microscopy was conducted in fetal sheep brain, and shows a similarly higher proportion of thin fibers in SWM and no difference in g-ratio (28). Interestingly, they also describe a higher proportion (8%) of thick (>8.5 µm) fibers in SWM, which we cannot confirm for the human brain. In the CC, a comparable light microscopic study found slightly larger fiber diameters (33). However, the authors report the radius of the equivalent circle instead of the short axis of an ellipse fitted on the structure, which likely explains the difference to the data presented here. The relationship of fiber diameter and g-ratio in the CC (Fig. 3d) integrates well with results from the human medullary pyramidal tract, which also consists of mostly long and thick fibers and is specialized on long-distance information transfer (34). Both our CC data and Keyserlingk & Schramm (34) show a large proportion of the larger fibers around 0.6 g-ratio, indicating it might be optimal for CNS long-range signal transmission. Our data also shows a higher number of fibers above g-ratio 0.6, which might have inaccuracies in the automated segmentation as a contributing factor. This, together with the more diverse g-ratio distribution (Fig. 3e), indicates that SWM likely features a larger variety of different fiber types as opposed to CC, and this results in a larger variety of g-ratios.

While we treat CC and SWM as single groups in this article, it is very likely that there are intra-regional variations of axonal properties within each group. In CC, Aboitiz and colleagues (29) describe axon diameters in subregions of the CC as decreasing from the genu to the posterior midbody, and then increasing from there to the splenium. In contrast, Riise & Pakkenberg (35) describe a linear upward trend of axon diameters from anterior to posterior regions of the human CC. Therefore, further studies examining local variations within CC are required.

In SWM, it is also possible that different specialized subregions exist. Yoshino and colleagues (36) describe SWM of ferrets with two distinct regions, where lower diameter fibers are very close (<50 µm) to the border of the cortex, highlighting local variability of SWM. Furthermore, they propose that U-fibers are more myelinated than deep white matter (37). We can not confirm these findings in humans. A case of potential local white matter plasticity in humans is within M1. In M1, each finger, wrist and elbow is represented in two areas of the primary motor cortex of humans: once in area 4a and once within area 4p (38,39). The representation in 4p is closely linked to the somatosensory perception and preferentially active during combined sensory and motor tasks, whereas area 4a is hypothesized to be connected more closely to the corticospinal tract and associative frontal areas (40–42). Therefore, if shorter SWM fibers are also thinner and less myelinated than long SWM fibers, SWM below 4a might have thicker and more myelinated fibers than 4p, where a larger proportion of fibers projects to neighboring somatosensory areas. Similarly, in the visual cortex, color-sensitive and ocular dominance columns in the primary visual cortex V1 have distinct connectivity patterns to thick, thin and pale stripes within V2, with potentially different structural properties of the different connecting fiber populations (12).

The conduction velocity of a fiber is a product of, among other factors, its axon diameter and myelination thickness. A correctly tuned conduction velocity is crucial for the coordination of activity across different brain regions through white matter. In the case of short association fibers directly connecting e.g. motor and somatosensory cortex are approximately 4 cm long. For such a fiber, with a mean conduction velocity of 3.1 m/s (see Fig. 3f), conducting a signal from motor to somatosensory cortex would take 13.8 milliseconds. Putting this into perspective with the typical chemical synaptic delay (∼1 ms), this makes a precisely tuned conduction velocity integral for synchronizing temporally coded neural activity in different parts of the brain.

The process of optimizing axon diameter and myelination thickness begins in the late prenatal period and continues until at least early adulthood (43). Even in the adult brain, white matter adapts to training stimuli over the course of weeks (5,44,45). Therefore, it is evident that white matter microstructure remains plastic in adults. It is currently unclear by which mechanism this precise tuning is regulated. While it is clear that activity can induce myelin plasticity, the molecular connection between these two phenomena remains poorly understood (46).

In humans, it is difficult to gather high quality electron microscopic data due to prolonged *post mortem* delay (20-40 hrs) before fixation and fixative perfusion not being a standard practice in a clinical pathology setting (47). Due to these issues, axons and myelin sheaths in electron microscopic data are frequently damaged, or, in the case of axons, degraded (Fig. 2a). This makes reliable and automated analysis of these data challenging. It is unknown if this degradation affects all fibers equally, regardless of axon diameter and myelin thickness. Therefore, a systematic bias related to the speed of degradation is possible. Furthermore, only brains of elderly individuals (60-78 years) were used for this study, which is likely to have an influence on the myelination state of white matter (48,49).

The strength of the current study is the very large number of analyzed axons, which was enabled by the use of deep learning algorithms for image analysis. These approaches may come with some limitations. Due to the ‘black-box’ functionality of deep learning approaches, we can not exclude possible systematic biases in the automated analysis of our electron microscopic data. For example, due to an imbalance in the amount of training data from CC and SWM regions, we might more reliably segment fibers in SWM that are more similar to CC fibers. While our validation efforts did not show concerning biases (see Fig. S1-S6,), it is possible that there are hidden effects. Furthermore, we ‘idealized’ obvious artifacts in the training data (e.g. partly broken down myelin sheaths), which is not typical and might contribute to systematic errors. If there are hidden effects, they are likely to be amplified for the measurement of the g-ratio, as the inaccuracy of two measurements (axon + outer fiber diameter) is multiplied in the process. The mean absolute error for the g-ratio (not including false positives and negatives) is 0.023 (CC) and 0.057 (SWM), see Fig. S5a,b.

In this study, we show a robust difference in white matter fiber composition of human SWM and CC. However, a variety of open questions remain. It is unclear how variable white matter structure is underneath neighboring cortical areas with different functions (e.g. primary motor areas 4a and 4p). Furthermore, there is still conflicting evidence about the differences of ultrastructure across CC subregions (29,35). We are currently investigating both of these questions. Another open question is a precise model of how myelination and fiber thickness is regulated and which factors contribute to the plasticity of white matter fibers. The presented methodological approach allows for analyzing large populations of fibers. It is therefore the method of choice to answer the above-mentioned questions.

The displayed data indicate that white matter structural variation between short and long fibers is present on the level of axon diameters and myelination thickness, but not in the (mean) g-ratio, albeit with a more complex g-ratio distribution in SWM than in CC. We hypothesize that this is due to a higher variety of different signaling distances (and therefore mixture of short and long fibers) present in SWM, which is manifested in a more diverse fiber composition. We argue that the axon and myelination thickness of a fiber results from a balance of complex interactions between various factors: The length of the fiber, physiological demands of the white matter tract it is part of, physical constraints of the tissue, and the local energy budget available.

With this study, we present the first large-scale analysis of white matter ultrastructure in the human brain, to our knowledge, proving a long-standing hypothesis in the field. While previous studies either relied on indirect measurement through MRI-based methods or on tissue from model animals with vastly different brain sizes (and therefore also vastly different connection distances), here we provide a comprehensive description of structural specialization in long and short white matter tracts. We show that fibers in short white matter tracts are thinner, less myelinated, but have the same mean g-ratio as fibers in longer tracts. Furthermore, we highlight how these structural specializations are integral to the correct tuning of signal conduction speeds. For future connectome studies in humans, we argue that it is essential to include structural information such as these, or information such as conduction velocity derived from these measures. White matter structure is finetuned to achieve optimal conduction velocity, energy efficiency, efficient volumetric packing of fibers, and precise conduction of temporally coded signals in the human brain.

## Methods

### Human samples

The brain samples used in this study (5 cases, 4 male, aged 61 (*post mortem* delay: 20h), 74 (*post mortem* delay: 24h), 78 (*post mortem* delay: 40h), 74 (*post mortem* delay: 24h), 60 (female, *post mortem* delay: 22h) years) were provided by the Paul Flechsig Institute – Center for Neuropathology and Brain Research at the Leipzig University Medical Faculty, the Department of Neuropathology Leipzig, and the Institute of Forensic Medicine at the University Medical Center Hamburg-Eppendorf, following ethical guidelines and with proper permissions. The entire procedure of case recruitment, acquisition of the patient’s personal data, the protocols and the informed consent forms, performing the autopsy, and handling the autopsy material have been approved by the responsible authorities (approval by the Ethics Committee of the University of Leipzig; approval #205/17-ek and approval #WF-74/16, by the Ethics Committee of the University Clinic Hamburg-Eppendorf. Neuropathological assessment revealed no signs of neurological diseases. All samples were immersion-fixed at 4°C in a solution of 3% formaldehyde and 1% glutaraldehyde in PBS buffer at pH 7.4 for at least 4 weeks before sectioning.

### Preparation of human samples

Samples were taken from SWM below primary motor, somatosensory, and primary and secondary visual cortex (see Fig. 1). For the motor- and somatosensory cortex, we identified the hand knob and the corresponding somatosensory region that is on the posterior side of the central sulcus.

CC samples were obtained from six regions (rostrum, genu, anterior midbody, midbody, isthmus, splenium) of the corpus callosum. CC samples were cut orthogonally to the long axis of the CC.

For electron microscopy, 70 µm thick vibratome sections were cut. At the same time, we also cut slices for histology, requiring 40 µm sections. The tissue block was embedded in low-melting agarose (2% agarose, heated, cooled down to 37°C, and set around the tissue block on a cold metal plate). Once solid, excess agarose was trimmed. The stage of the vibratome (HM 650 V, Thermo Scientific) was coated with superglue, the block placed on top, and weighed down. After the glue set, the stage was installed in the vibratome, with the surrounding bath filled with PBS. The cutting process involved two steps: a coarse trim of 125–250 µm, then fine cuts for the 70 µm sections for electron microscopy and 40 µm for histology. The cutting speed was 1–1.4 mm/s, with a frequency of 70 Hz and a knife amplitude of 0.8 mm. The thicker sections for electron microscopy were stored in 0.2M cacodylate buffer, while the thinner histology sections were kept in PBS-azide.

### Nissl staining

Vibratome sections of 40 µm thickness were used for Nissl staining. Sections were mounted on glass slides and air dried for 10 minutes at 40°C, then rehydrated and rinsed in distilled water for three minutes. The sections were then moved through a graded series of ethanol (70%/85%/95% ethanol for 5 minutes each), and then kept in 95% ethanol for 30 minutes. Then, they are moved through a descending alcohol series (95%/85%/70% ethanol) for 5 minutes each. Following this, the sections were thoroughly rinsed with distilled water several times.

The samples were stained for ten minutes with an aqueous solution of 0,1% cresyl violet and quickly rinsed with distilled water. The sections then underwent differentiation through a series of ethanol concentrations (70%, 85%, two rounds of 95%, and finally 100%) for approximately 2-5 minutes under visual control. After achieving the desired level of differentiation, the sections were rinsed twice with toluene and then mounted with Entellan in toluene under a fume hood.

### Transmission Electron Microscopy - Sample Preparation

For TEM imaging 70 µm thick vibratome sections were prepared using a Microm HM 650 V (Thermo Scientific, Walldorf, Germany). Small sections targeting the superficial white matter, including adjacent grey and deep white matter were cut out and stored at 4°C in 0.2M cacodylate buffer (4.28% Na-cacodylate, 1.2% 1N HCl) until further processing. They were contrasted in 1% osmium tetroxide in cacodylate buffer for 1 hour at room temperature on a laboratory shaker (Heidolph Rotamax, 70 rpm), followed by two rinses in cacodylate buffer, each for at least 1 hour, and then stored overnight at 4 °C. Dehydration was carried out through a series of acetone solutions (30 % for 15 min, 50 % for 30 min, 70 % for rinsing) with a final contrast enhancement step using 1% uranyl acetate in 70% acetone for 45 min, protected from light, and rinsed in 90% acetone for 30 min on a shaker (70 rpm) and then two times 100% anhydrous acetone for 30 minutes each. The samples were embedded in Durcupan^TM^ araldite resin (Sigma-Aldrich Chemie GmbH, Steinheim) by sandwiching them between polyethylene terephthalate foils on glass slides. Once cured, the backing foil was removed for bonding on a Durcupan block and the remaining foil was peeled away. Initial 0.5 µm semithin-sections were stained with toluidine blue and imaged using the AxioScan Z1 slide scanner for orientation. For large field of view transmission electron microscopy, 50 nm sections were prepared with a Leica Reichert Ultracut S and mounted on slot grids (SF 162 N5 Provac, type 1x2 mm^2^) coated with a 10–20 nm formvar film.

### Transmission Electron Microscopy - Imaging

Digital electron micrographs were acquired at 80 kV using the LEO 912 EM OMEGA (Carl Zeiss, Oberkochen, Germany) equipped with an on-axis YAG scintillator and a 2k x 2k CCD 16-bit camera with ImageSP software (TRS, Moorenweis, Germany). Large area field of views (75 µm x 750 µm) were achieved by automatic tile scanning with 10% overlap and a pixel resolution of 4,32 nm. The resulting images were stitched using ZEISS ZEN (version 3.9). The stitched images were then manually cropped to exclude border areas of the imaging slide with no tissue on it and physically damaged areas of the tissue.

### Transmission Electron Microscopy analysis - Segmentation

We labelled ∼16000 cells manually as training data and trained a DenseNet (31,50)using Uni-EM (51). For inference, we downsampled the raw data by the factor of 16 (4 per axis) and applied a local contrast enhancement (CLAHE, Contrast Limited Adaptive Histogram Equalization). Prediction was conducted via the inference interface of Uni-EM, using a receptive field of 2048x2048. For the post processing after prediction, the semantic segmentation mask (Fig. 2b) was thresholded and corrected with binary operations and filtered for unrealistic fibers (e.g. g-ratio >1, unrealistically large or small fibers).

All custom python code is available at github.com/PhilipRuthig/ShortvsLongfibers/. In brief, we preprocessed the data with a CLAHE filter to enhance local contrast (Fig. 2a). These pictures were then predicted by the trained DenseNet to yield a semantic segmentation (Fig. 2b). The semantic segmentation was post processed to separate different instances of axon + myelination pairs (Fig. 2c) and separate ‘kissing’ instances by watershedding from each center of a segmented fiber.

The measures taken from the segmented binary structures were taken on ellipses with the same normalized second moment as each structure. The measurements taken from each structure include: minor & major axis, eccentricity, orientation, and g-ratio. For this study, we defined the g-ratio as the ratio between the minor axis of the ellipse fitted on the axon divided by the minor axis of the ellipse fitted on the myelination, as these are the most reliable measures according to our validation (see Fig. S1,S2).

All custom python scripts used in this work are based primarily on numpy (52), scikit-image (53), and scipy (54). All plots except Fig. S7, S8 were generated with Matplotlib (55) and Pylustrator (56).

### Transmission Electron Microscopy analysis - Validation

For the validation of our segmentation pipeline, we trained a validation model on a subset of randomly chosen labeled training images and validated its prediction against the last image not inside the subset. The segmentation pipeline reached a IoU scores of 0.86/0.79 for axons and myelination in CC data, and 0.78/0.58 for SWM data. We compared population-level differences of the validation and prediction of the same data (Fig. S1, S2) and compared instance-level differences by matching instances with >40% identity in prediction and validation data. All instances that had no corresponding fiber in the prediction or validation data were treated as false positives or negatives, respectively. The remaining fibers were plotted against each other in a paired way (Fig. S3, S4), and Bland-Altman plots were created analogous to Bland & Altman (57,58). Furthermore, we show the individual error for every reported measure per fiber instance (Fig. S5).

Since, in SWM, the cutting angle is more likely to be oblique, we also added a control for cutting angle to our data. We cut half of our SWM samples in parallel to the main fiber direction described by blunt dissections (1), and the other half orthogonally. We show the resulting axon diameters in Fig. S6.

### Bayesian modeling

We test-fitted a set of 18 probability distributions on data measured on manually labeled data, for which the GEV gave the best fit on axon diameters (data not shown). Therefore, we chose to employ the same distribution in this publication. It is defined by a location parameter *µ*, scale parameter δ and a shape (‘tailedness’) parameter ξ according to Coles (59).

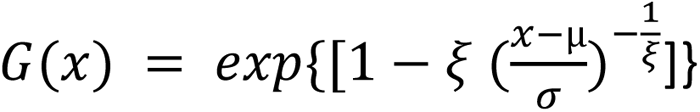

For the Bayesian statistics used in this study, we used PyMC version 5.10.0 and PyMC experimental 0.0.15. Modelling-related visualizations (Fig. S6,7) were generated using Arviz 0.16.1 (60). For prior information, we chose the same weakly informative priors (based on a GEV fit to the manually labelled training data for our deep learning model) for SWM and CC data. The models were designed analogous to the PyMC case study “Generalized Extreme Value Distribution” by Colin Caprani (61,62).

### Conduction velocity estimation

We estimated the conduction velocity *V* of the segmented fibers based axon diameter *d* and g-ratio *g* according to Rushton (18), with a modified exponent for the square root (0.68 instead of 0.5) as suggested by Schmidt & Knösche (32). Furthermore, as Rushton’s formula only returns relative conduction velocities, we multiplied the resulting velocity with a factor of 7.5 to match the experimental results displayed in Schmidt & Knösche (32):

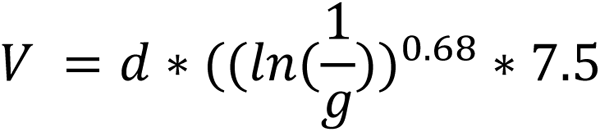

## Code Availability

All of the code used for the analyses in this study is publicly available on Github (<u>github.com/PhilipRuthig/ShortvsLongfibers/</u>). The repository also contains sample image data of CC and SWM of every step in the pipeline. The deep learning model and accompanying training data will be published at a later date, but are available upon request.

## Data Availability

All of the raw and processed data is available upon request. All text-based data (csv files with axon & outer fiber diameters, g-ratio measurements) and scripts that process these data are available at <u>github.com/PhilipRuthig/ShortvsLongfibers/</u>.

## Contributions

M.M., E.K., S.M., N.W. conceptualized, designed, supervised, and acquired funding for the study. P.R. and D.E.v.d.P wrote the manuscript, with review and editing by M.M., E.K., S.M., and N.W. The data were generated by Maria M., P.R., and D.E.v.d.P., K.R., and T.R.. Manually labeled training data was generated by Maria M., P.R., and D.E.v.d.P. All analyses and accompanying code were designed and written by P.R., with support from S.M. for the validation. P.R. designed the visualizations. Project administration was done by M.M.

## Acknowledgements

We sincerely thank Laurin Mordhorst, Torsten Bullmann, Thomas Knösche, Maëlig Chauvel, Gesine Müller, and Nico Scherf for helpful discussions regarding the analyses conducted in this manuscript. We also thank Ruth Stassart for her comments on earlier versions of the manuscript. Further, this work was supported by the German Research Foundation (DFG Priority Program 2041 ‘Computational Connectomics’ MO 2249/3-1, MO 2249/3-2, KI 1337/2-2, WE 5046/4-2). Figure 1a was created using biorender.com. SM was additionally supported by the DFG Emmy Noether program MO 2397/4-1 and MO 2397/4-2 and the ERC (Acronym: MRStain, Grant agreement ID: 101089218, DOI: 10.3030/101089218). Views and opinions expressed are however those of the authors only and do not necessarily reflect those of the European Union or the European Research Council Executive Agency. Neither the European Union nor the granting authority can be held responsible for them.

## Supplementary Materials

**Fig. S1:**
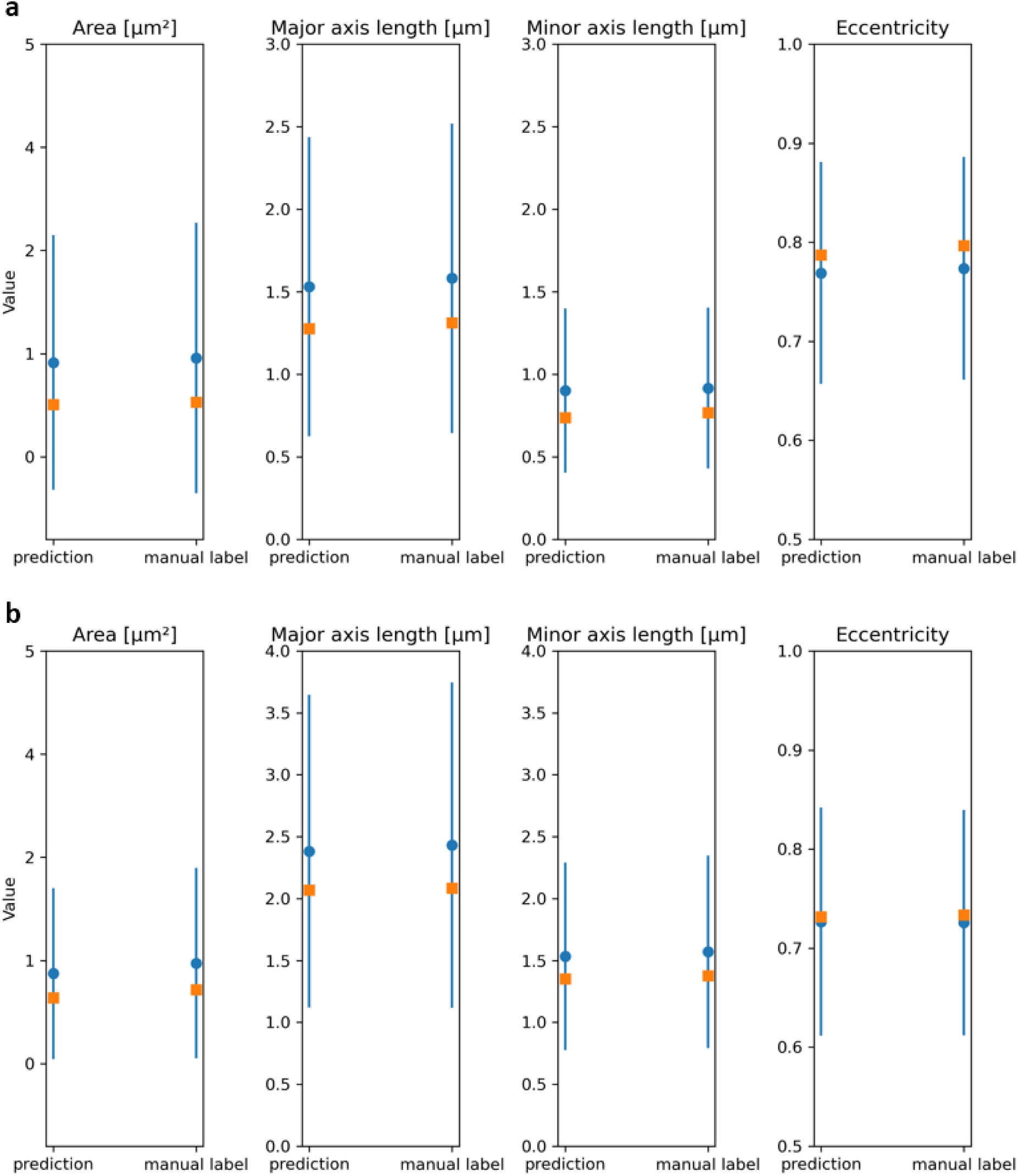
Population-level measures of leave-one-out validation in CC. This plot shows population-level differences in the training data (manual label) and our prediction, showing only minor systematic biases in the data. All plots show mean ± sd (blue) and median (orange) of area, major & minor axis length and eccentricity of CC axons (**a**), CC outer fiber measures (**b**).

**Fig. S2:**
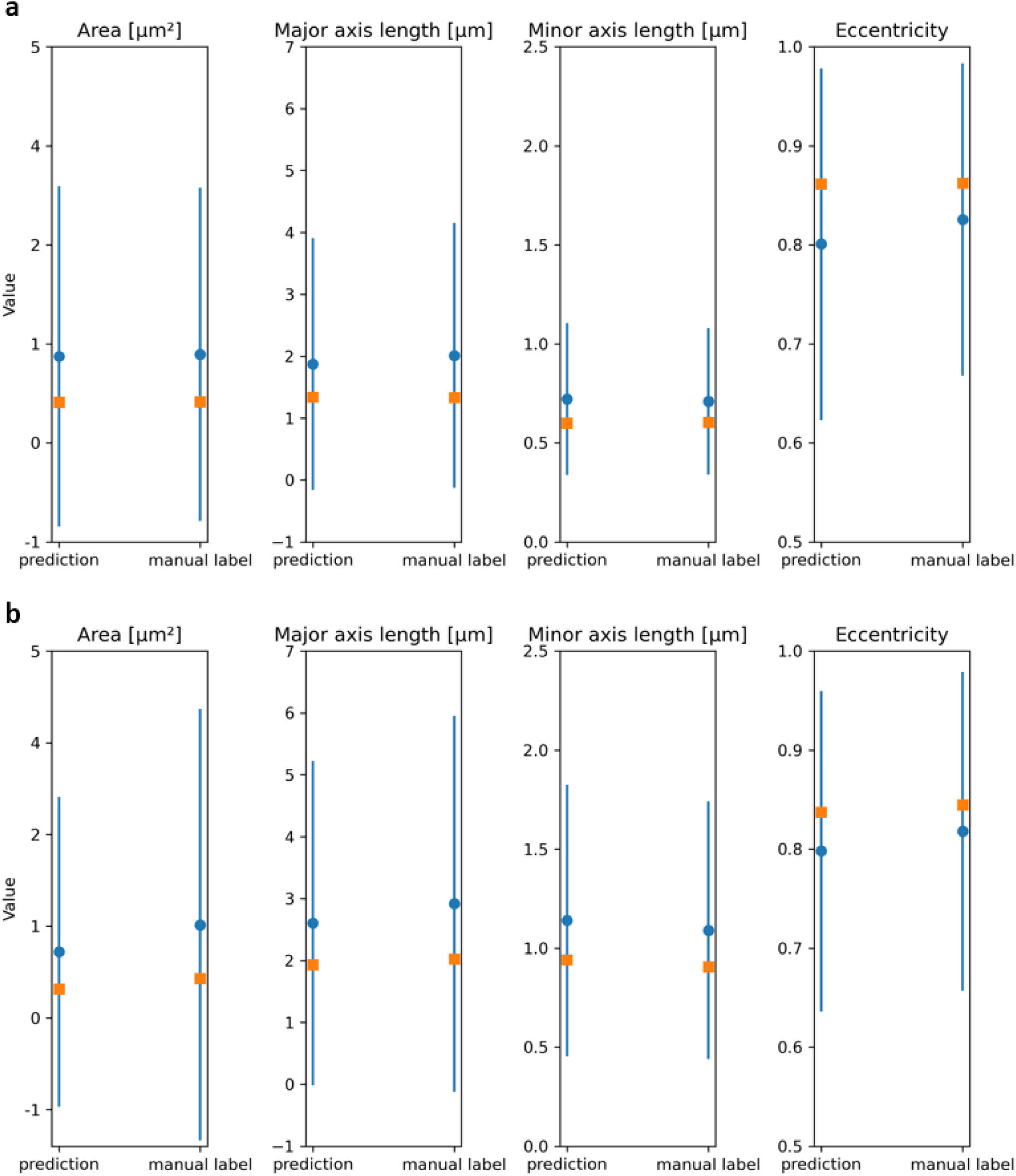
Population-level measures of leave-one-out validation in SWM. This plot shows population-level differences in the training data (manual label) and our prediction, showing only minor systematic biases in the data. All plots show mean ± sd (blue) and median (orange) of area, major & minor axis length (in µm) and eccentricity of SWM axons (**a**), SWM outer fiber measures (**b**).

**Fig. S3:**
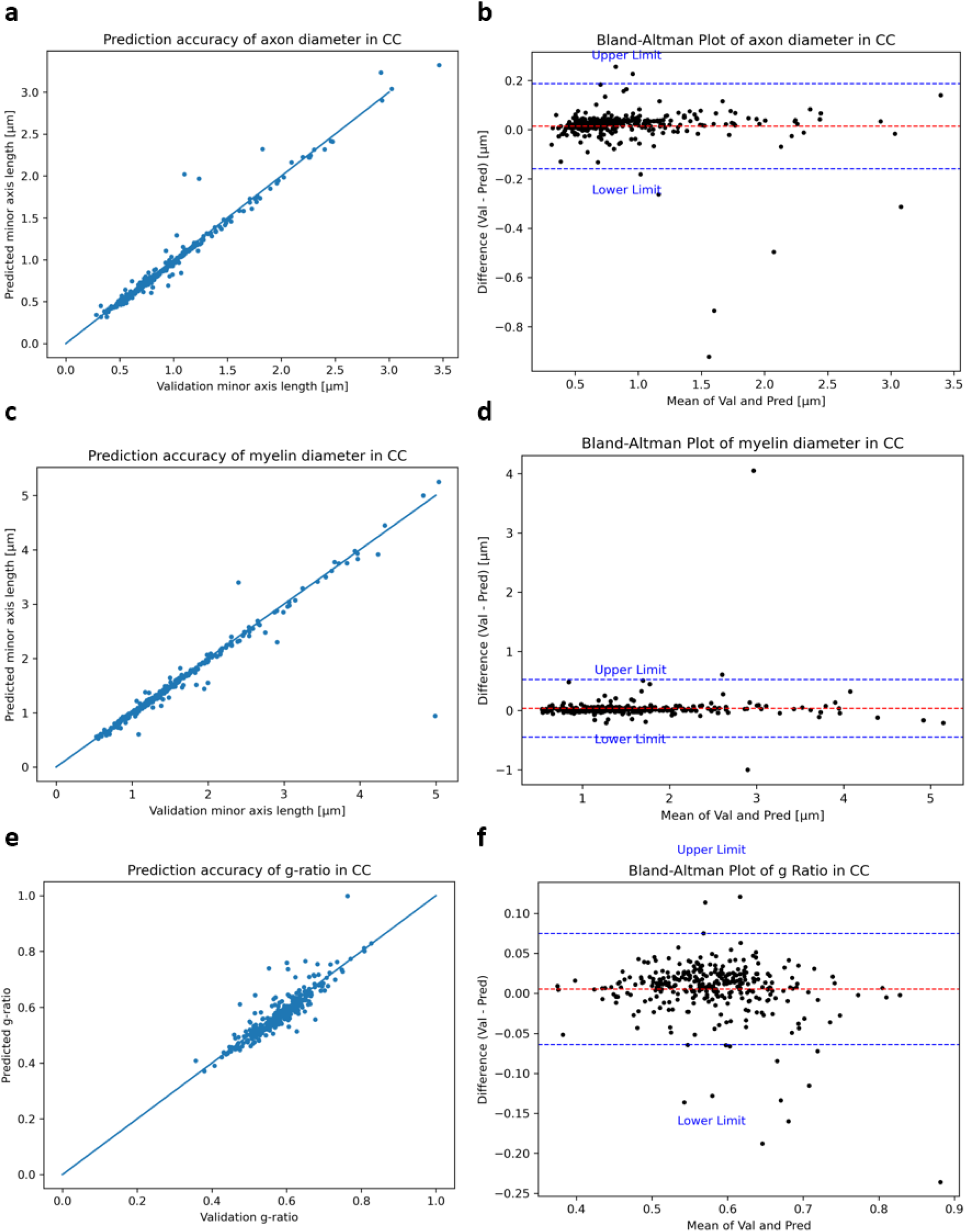
Individually paired data of leave-one-out validation of segmentation algorithm in CC. To validate the outputs from automated segmentation, we validated them against manually labeled data identical to the training data (but not part of the training dataset). Summaries on population level are shown in S2. Here, all plots show paired data, with each point being a single fiber in the prediction and validation image having more than 40% overlap with the corresponding validation fiber. Plotted lines show theoretical perfect correlation. Plots show correlation of predicted measures of each structure with the validation data (left column) and Bland-Altman plots (right column) of CC axon diameter (**a**,**b**), CC outer fiber diameter (**c**,**d**), CC g-ratio (**e**,**f**).

**Fig. S4:**
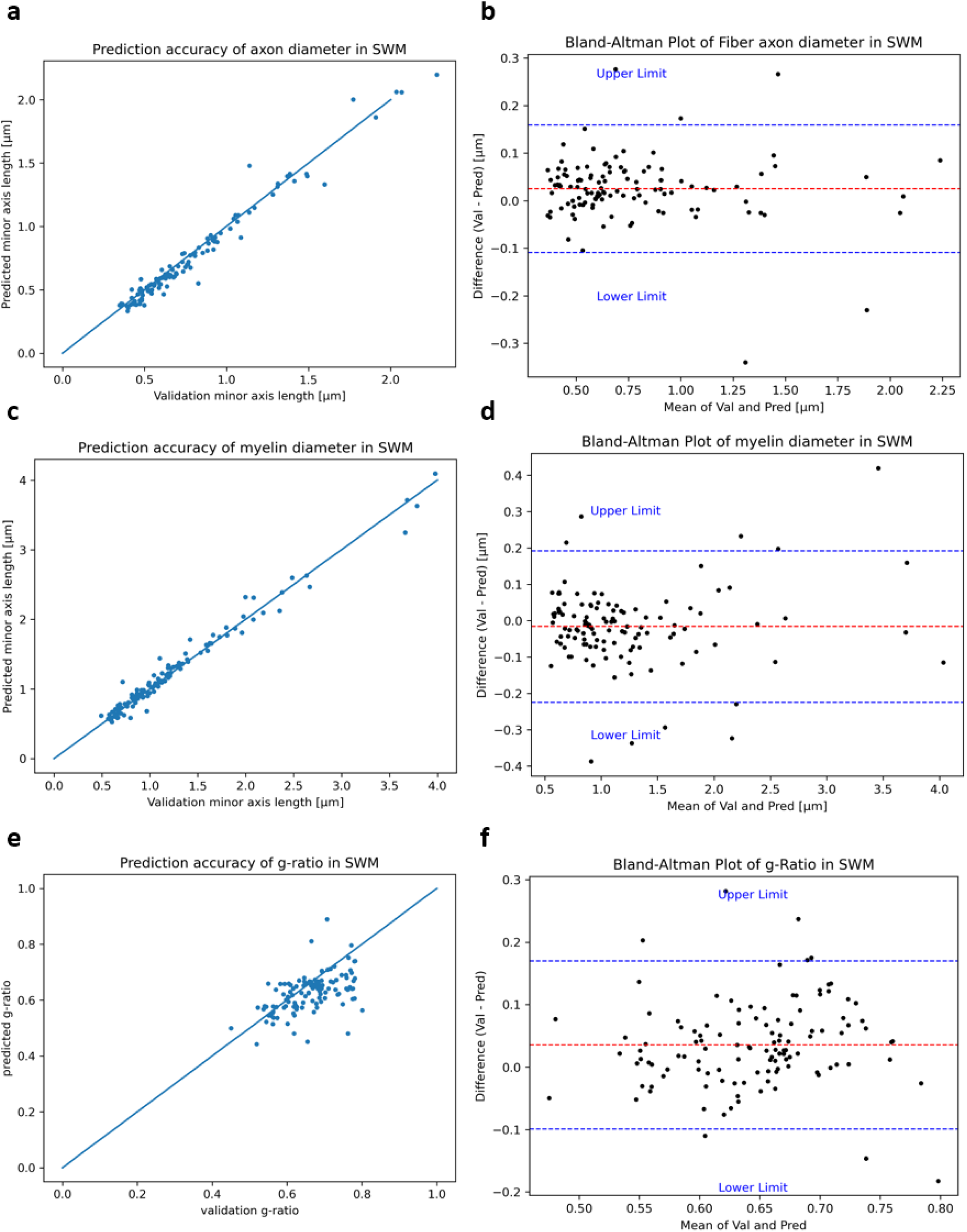
Individually paired data of leave-one-out validation of segmentation algorithm in SWM. To validate the outputs from automated segmentation, we validated them against manually labeled data identical to the training data (but not part of the training dataset). Summaries on population level are shown in S2. Here, all plots show paired data, with each point being a single fiber in the prediction and validation image having more than 40% overlap with the corresponding validation fiber. Plotted lines show theoretical perfect correlation. Plots show correlation of predicted measures of each structure with the validation data (left column) and Bland-Altman plots (right column) of SWM axon diameter (**a**,**b**), SWM outer fiber diameter (**c,d**), SWM g-ratio (**e,f**).

**Fig. S5:**
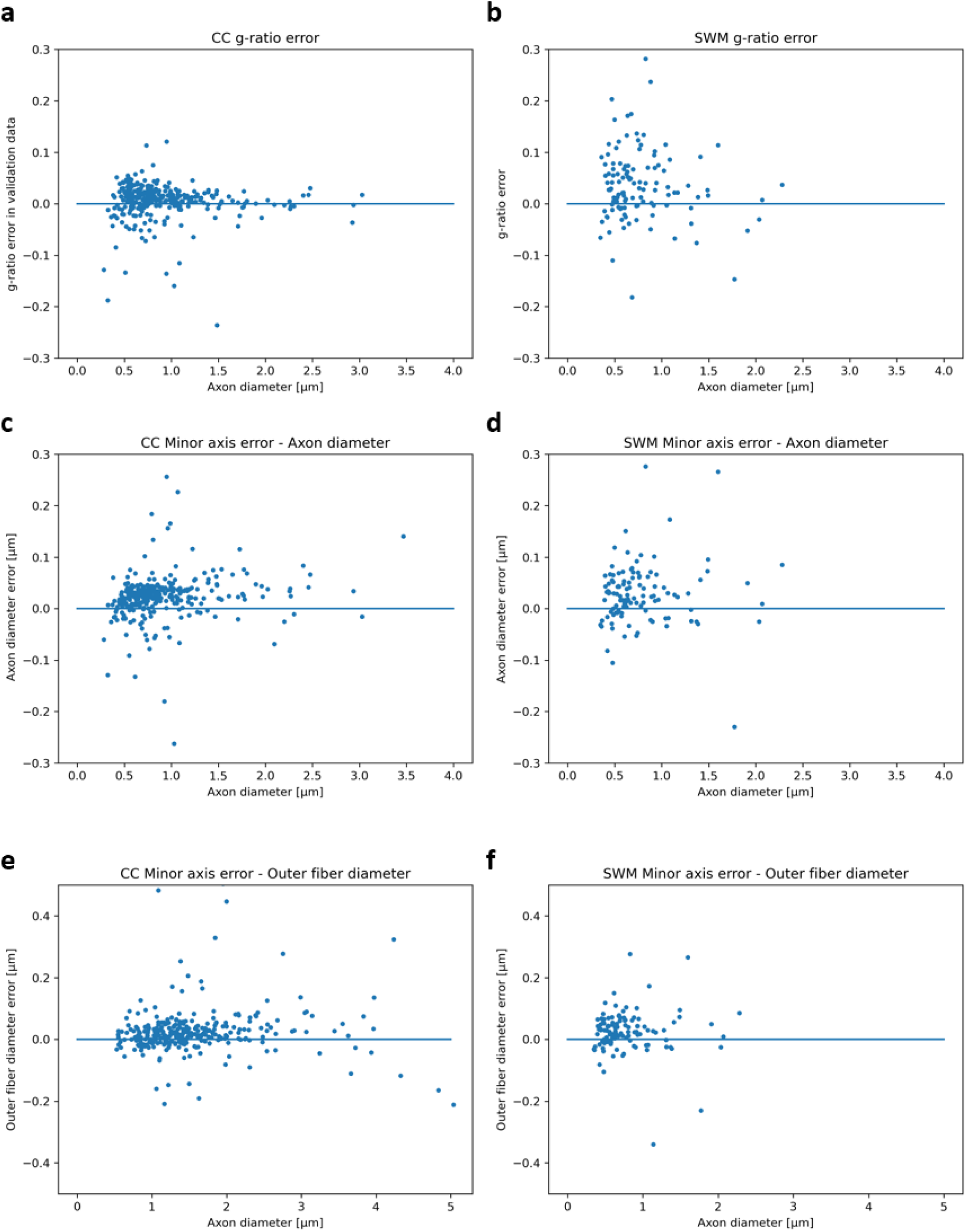
Measurement error for paired data, measured against axon diameter. Plots show the axon diameter of the validation instance plotted against the measurement error (delta of validation - prediction value) of that instance. These paired error values are shown for g-ratio in CC (**a**) and SWM (**b**), axon diameter in CC (**c**) and SWM (**d**), and outer fiber diameter in CC (**e**) and SWM (**f**).

**Fig. S6:**
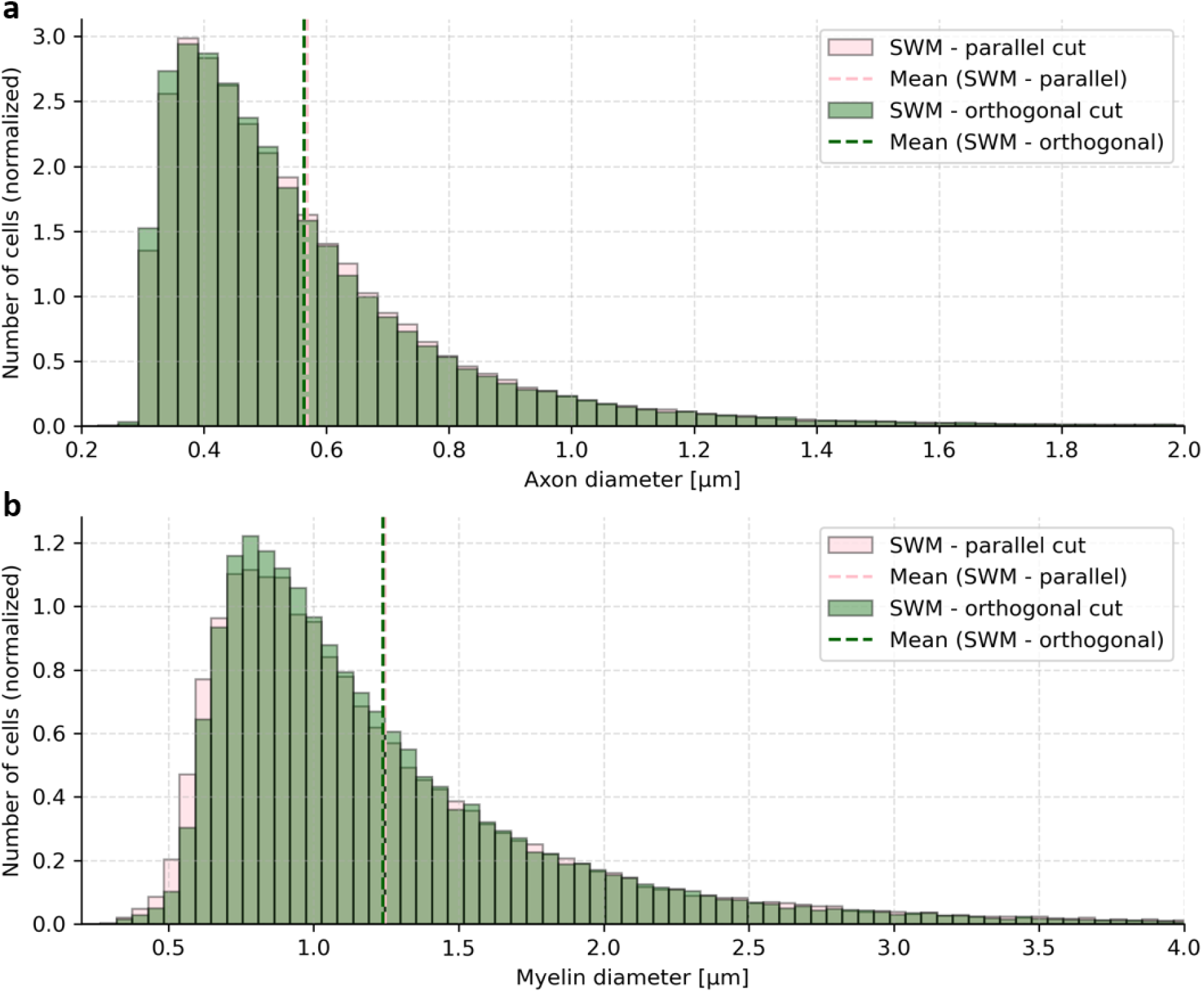
Raw axon and outer fiber diameters for parallel and orthogonally cut SWM. **a,b,** Axon diameters (**a**) and outer fiber diameters (**b**) of SWM data, split according to cutting angle. A portion of the data was cut orthogonal to the main short association fiber direction, and a portion was cut in parallel to the main short association fiber direction. Two example slices for the parallel cut can be seen in Fig. S1 c,d. Orthogonal cuts are not shown.

**Fig. S7:**
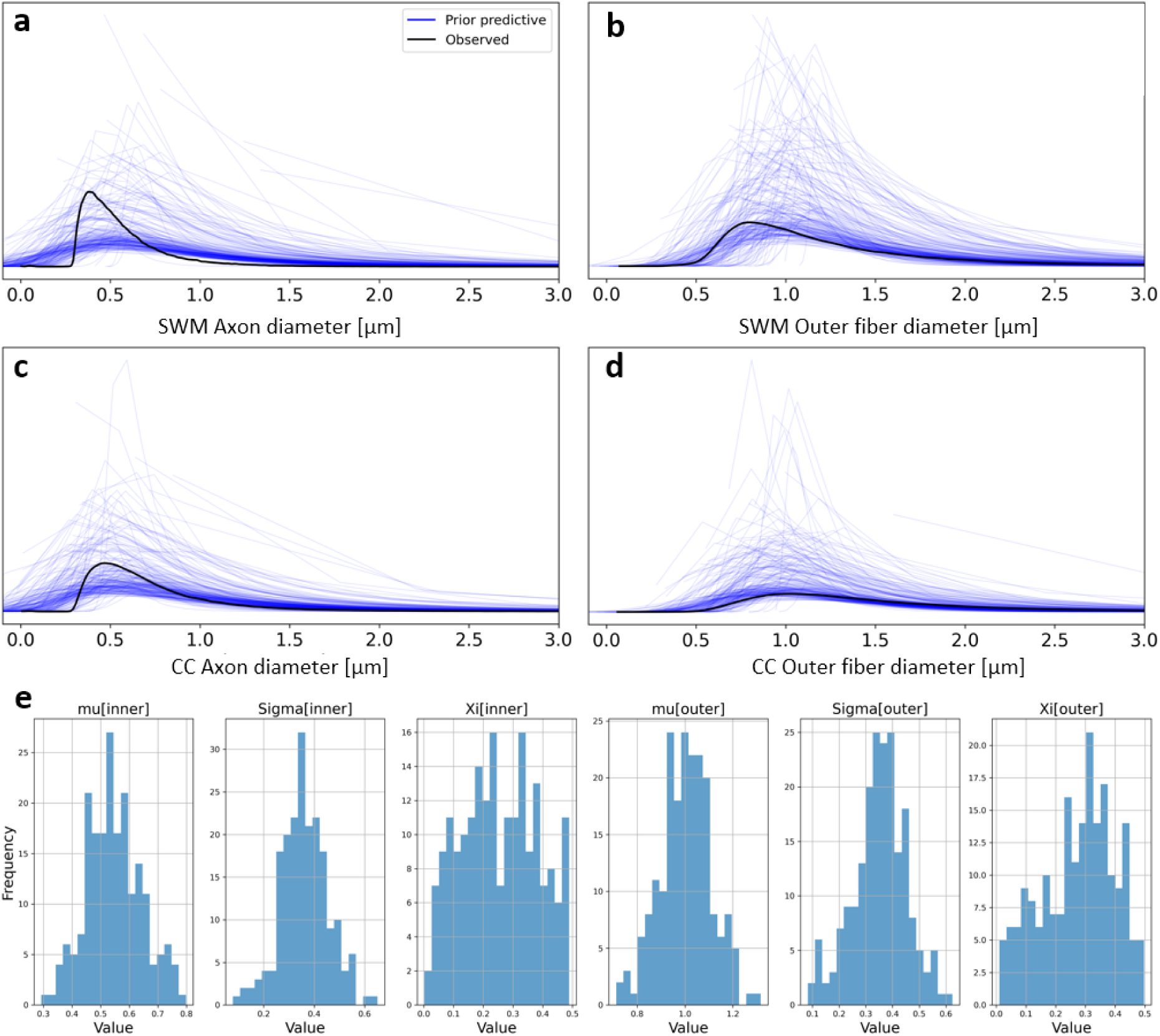
Prior predictive sampling for fitting a GEV distribution to CC and SWM data. **a,b,c,d** show prior predictive GEVs for SWM axon diameter (**a**), SWM outer fiber diameter (**b**), CC axon diameter (**c**), and CC outer fiber diameter (**d**). **e,** Histograms of 200 Prior samples for each of the parameters.

**Fig. S8:**
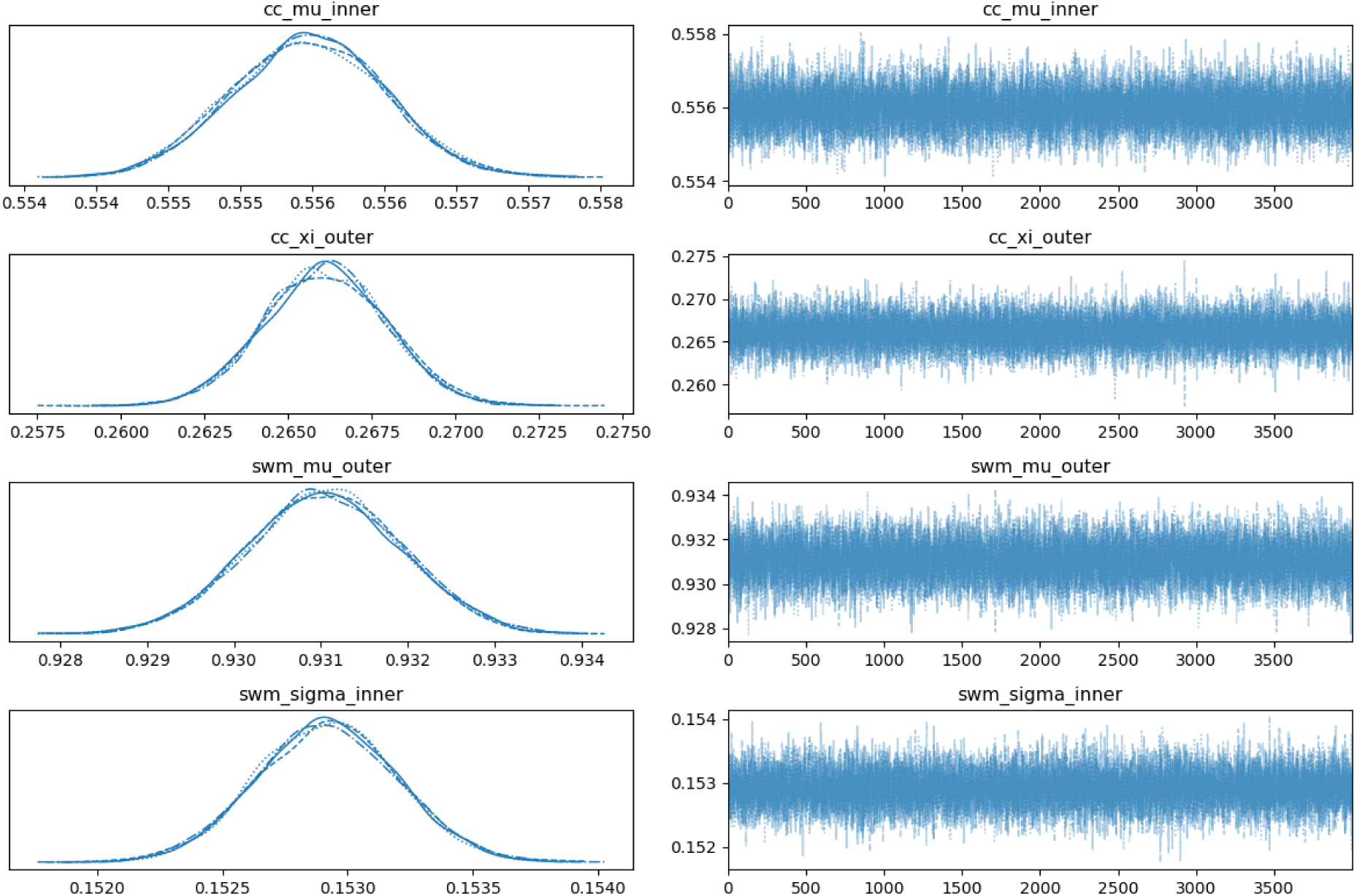
Sample traceplots from MCMC sampling. Left column shows histograms of all posterior samples for a set of 4 parameters. Different line styles represent 4 different chains. Right column shows trace plots of the Markov Chain Monte Carlo sampling procedure for a single chain each.

